# Modification of primary amines to higher order amines reduces in vivo hematological and immunotoxicity of cationic nanocarriers through TLR4 and complement pathways

**DOI:** 10.1101/647305

**Authors:** Randall Toy, Pallab Pradhan, Vijayeetha Ramesh, Nelson C. Di Paolo, Blake Lash, Jiaying Liu, Emmeline L. Blanchard, Philip J. Santangelo, Dmitry M. Shayakhmetov, Krishnendu Roy

## Abstract

For decades, cationic polymer nanoparticles have been investigated for nucleic acid delivery. Despite promising in vitro transfection results, most formulations have failed to translate into the clinic due to significant *in vivo* toxicity – especially when delivered intravenously. To address this significant problem, we investigated the detailed mechanisms that govern the complex *in vivo* systemic toxicity response to common polymeric nanoparticles. We determined that the toxicity response is material dependent. For branched polyethylenimine (bPEI) nanoparticles – toxicity is a function of multiple pathophysiological responses – triggering of innate immune sensors, induction of hepatic toxicity, and significant alteration of hematological properties. In contrast, for chitosan-based nanoparticles – systemic toxicity is primarily driven through innate immune activation. We further identified that modification of primary amines to secondary and tertiary amines using the small molecule imidazole-acetic-acid (IAA) ameliorates *in vivo* toxicity from both nanocarriers by different, material-specific mechanisms related to Toll-like receptor 4 activation (for bPEI) and complement activation driven neutrophil infiltration (for chitosan), respectively. Our results provide a detailed roadmap for evaluating *in vivo* toxicity of nanocarriers and identifies potential opportunities to reduce toxicity for eventual clinical translation.

**Graphical Abstract:** 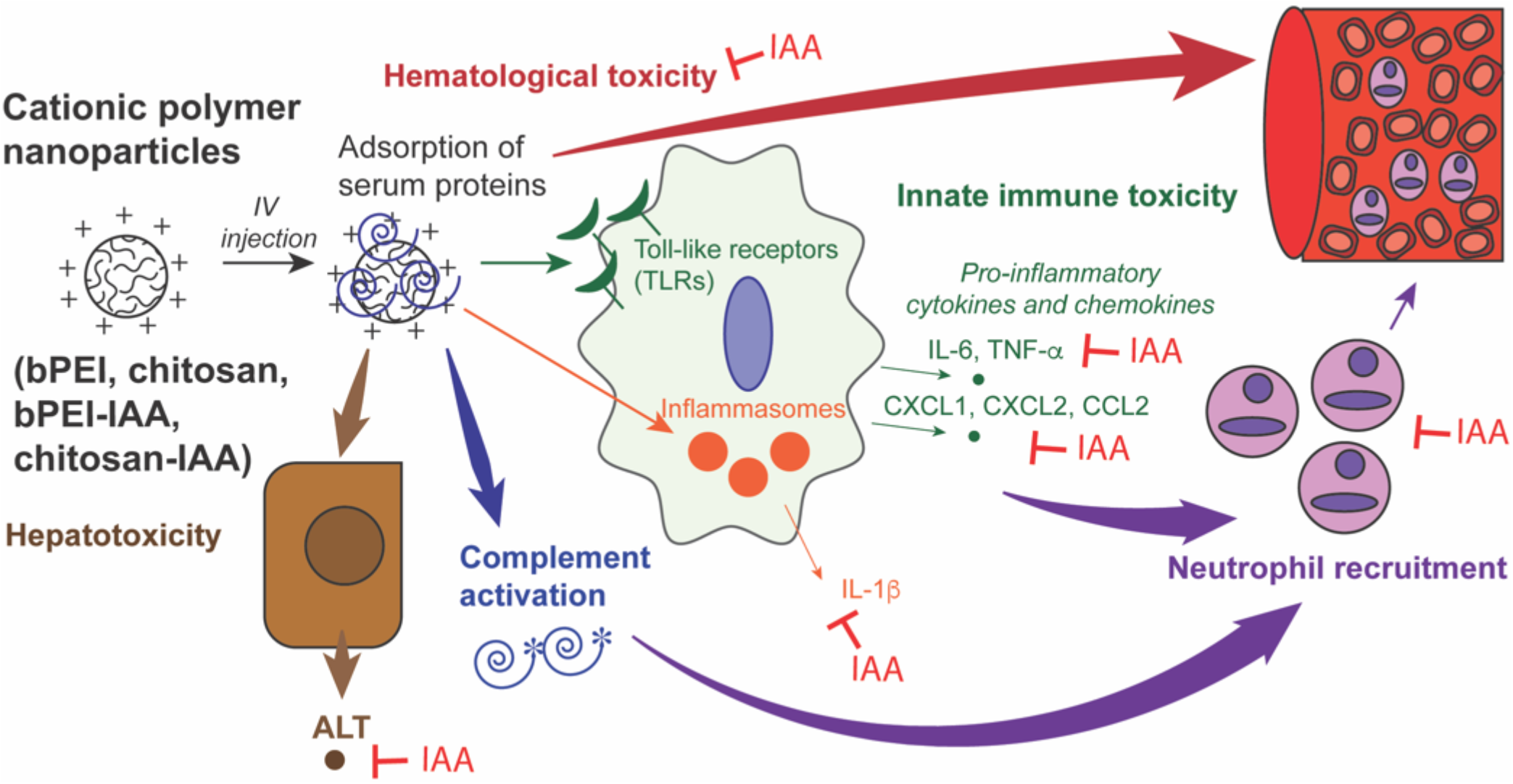

## INTRODUCTION

A wide variety of nanoparticles have been developed over the past decades to facilitate the delivery of nucleic acid-based therapies, such as plasmid DNA, oligonucleotides, short-interfering RNAs (siRNA), micro RNA (miRNA) and messenger RNA (mRNA).[1, 2] To form nanoparticles, anionic nucleic acids are often complexed with cationic polymers or cationic lipids – a process that results in nanoparticles with efficient nucleic acid loading and favorable physical characteristics for intracellular delivery of the cargo in vitro.[3],[4] Despite several desirable attributes, cationic nanoparticles induce significant *in vivo* toxicity, especially when delivered systemically, which has prevented their clinical translation even after decades of research. For lipid-based particles, toxicity profiles have been improved by using ionizable and neutral lipids with other excipients. In fact, most current clinical trials for systemic delivery using nanoparticles use lipid-based particles.[5–7] Meanwhile, a whole class of materials (cationic polymers), despite demonstrating promising in vitro efficacy results for decades, has not advanced into the clinic. In some cases nanoparticles consisting of modified cyclodextrin or protamine were evaluated in Phase I clinical trials, but further studies were not pursued.[8, 9]

An ideal nucleic acid delivery agent must ensure a balance between low toxicity and high therapeutic efficacy in vivo – two attributes that have been difficult to achieve simultaneously using cationic polymer nanoparticles.[10] Chemists and material scientists have empirically optimized physicochemical properties (e.g. size, surface functional groups, charge) to improve nanoparticle safety and transfection efficiency [11–17] These studies identified formulations with lower *in vitro* toxicity, and in some cases, the potential for *in vivo* efficacy in animal models, but detailed studies on the relationship between nanoparticle design and mechanisms of *in vivo* systemic toxicity have been scarce or very limited in scope.[18] A widely studied modification to reduce systemic toxicity of nanoparticles is the addition of polyethylene glycol (PEG) to the formulations. A major drawback of PEG-modified formulations, however, is that anti-PEG antibodies can be generated, which activate the complement cascade and further decrease therapeutic efficacy.[19–22] This shortcoming of PEG, which could not be predicted *in vitro*, highlight the importance of studies that investigate the detailed *in vivo* mechanisms by which nanoparticles induce systemic toxicity.

Accurate prediction and subsequent minimization of *in vivo* systemic toxicity requires a critical and comprehensive understanding of how nanoparticles interact with a variety of biological components after in vivo delivery, e.g., with serum proteins, blood and immune cells, and cells in various tissues like the liver, lung, and spleen. It is also critical to understand how they distribute to organs and in specific cell-types to trigger the innate immune response. In this study, we identify the distinct contributors to the *in vivo* toxicity responses induced by two widely-reported cationic nanoparticles for nucleic acid delivery – branched polyethyleneimine (bPEI) and chitosan. We found that the mechanism of in vivo systemic toxicity is a combination of innate immune responses, complement reactions, and hepatic and hematological toxicities – and the relative contribution of these differs with the type of polymer. We then compared *in vivo* systemic toxicity to these nanoparticles after modification with the small molecule imidazole-acetic-acid (IAA), which modifies primary-amines on the polymers and introduces secondary and tertiary amines. We hypothesized that such modification would reduce innate immune activation by downregulating interactions with the Toll-like receptor 4 (TLR4). While we observed reduced TLR4-mediated systemic toxicity with the IAA-modified nanoparticles, we also observe a significant impact on other measures of the systemic toxicity response – which include *in vivo* hepatotoxicity, complement activation, and changes in hematological properties. Investigation *of in* vivo systemic toxicity in TLR4 knockout mice confirmed that the mechanisms of toxicity are not completely TLR4-driven. Our results indicate that alleviation of in vivo toxicity by IAA is material-dependent and is driven by unique physiological pathways.

These observations provide valuable insights on the *in vivo* immune response to cationic nanoparticles for nucleic acid delivery. They also established a detailed roadmap to better understand the origins of *in vivo* systemic toxicity of nanoparticles. The insights generated could allow rational design of improved cationic polymer formulations that can achieve optimal balance between toxicity and efficacy *in vivo* – thereby enabling their clinical translation.

## MATERIALS AND METHODS

### Animals

All animal experiments were conducted in accordance to approved IACUC protocols at the Georgia Institute of Technology and Emory University (Shayakhmetov Lab). Female, 8-12 weeks old C57/Bl6 mice (Jackson Labs, Bar Harbor, ME) were used for all wild-type toxicity and mRNA transfection studies. Female, 8-12 weeks old TLR4 knockout mice (Jackson Labs, Bar Harbor, ME) were used in knockout studies. Euthanasia was performed with CO2 and confirmed by cervical dislocation.

### Synthesis and characterization of bPEI and chitosan nanoparticles

Unmodified bPEI nanoparticles (BPEI-TPP) and IAA-modified bPEI nanoparticles (bPEI-IAA-TPP) were produced by mixing solutions of bPEI or bPEI-IAA with sodium tripolyphosphate in deionized water. The mass ratio of bPEI or bPEI-IAA to TPP was 3:2. Unmodified chitosan nanoparticles (chitosan-TPP) and IAA-modified chitosan nanoparticles were produced by mixing solutions of chitosan or chitosan-IAA with sodium tripolyphosphate in sodium acetate buffer (pH= 4.5). The mass ratio of chitosan or chitosan-IAA to TPP was 5:1. Following vortexing, the nanoparticle formulations were then put onto a rotator at room temperature for 30 minutes. After rotating, nanoparticles were centrifuged at 4000 g for 20 minutes in Amicon centrifuge filters (100 kD MWCO) and resuspended in 1 mL PBS. The size and zeta potential of the particles were measured using a Malvern Zetasizer Nano ZS. For fluorescent labeling, bPEI and chitosan nanoparticles were reacted with Vivotag-645 for 1 hour on a rotator in the dark. Unreacted dye was removed by centrifugation at 4000 g with Amicon centrifuge filters (100 kD MWCO). Fluorescent nanoparticles were re-suspended in 1 mL PBS prior to use in experiments.

### *In vivo* nanoparticle-induced acute systemic toxicity

Mice were administered an 8 mg chitosan/kg body weight or 3 mg bPEI/kg body weight IV dose of nanoparticles. At specified endpoints, mice were sacrificed and blood or spleens were collected. Blood was stored at 4 C overnight and centrifuged at 12,000 g for 5 minutes to isolate serum. The IL-6, IL-1β, and TNFα ELISAs were performed on serum samples using Ready-Set-Go kits (eBioscience, San Diego, CA). ALT assays (Biovision, Milpitas, CA) were performed for serum samples. Histamine, complement C3a, and complement C5a levels in serum were quantified by ELISA (Histamine kit: Enzo Life Sciences, Farmingdale, NY; Complement C3a kit: Quidel, San Diego, CA; Complement C5a kit: Sigma-Aldrich, St. Louis, MO). Spleen tissues were homogenized with a FastPrep-24 instrument in RIPA buffer supplemented with EGTA, proteinase inhibitors, and Triton X (1%). Lysates were frozen at – 70 C and thawed. Cell debris was pelleted out by centrifugation at 10,000 g for 5 minutes. Blots of spleen lysates were prepared with a Proteome Profiler Mouse Cytokine Array Kit (Panel A, R&D Systems, Minneapolis, MN) using an established protocol.[23] After blot development, chemiluminescent imaging was performed using an Amersham Imager (GE Healthcare, UK).

### Complete blood count (CBC) analysis

Mice were administered a 3 mg bPEI/kg or 12 mg chitosan/kg body weight IV dose of nanoparticles. After 30 minutes or 150 minutes, mice were sacrificed and blood was collected in vials with a lithium heparin gel as anticoagulant. Hematocrit levels, hemoglobin concentration, and neutrophil fractions of white blood cells were measured in a Sysmex XN-L Automated Hematology Analyzer.

### Proximity ligation assay

RAW 264.7 macrophages were seeded in 96 well plates at a density of 30,000 cells/well. Cells were treated with 1.8 μg of nanoparticles for 30 minutes. At 30 minutes, cells were washed three times with PBS, fixed with 1% paraformaldehyde in PBS for 10 minutes at room temperature, and permeabilized with 0.2% T ritonX-100 for 5 minutes at room temperature. PLA was then performed between TLR4 (Novus NB100-56581, 1:50 dilution) and TIRAP (Novus NB300-990, 1:100 dilution) and analyzed via flow cytometry using a DuoLink PLA flow cytometry kit (Sigma Aldrich, St. Louis, MO). Samples were analyzed using a BD Accuri C6 Benchtop Cytometer and median fluorescence intensity was reported.

### *In vitro* nanoparticle-induced acute toxicity in BMDCs

Bone marrow cells were harvested from C57 Bl/6 mice. The cells were differentiated into dendritic cells using mouse GM-CSF (20 ng/mL; Peprotech, Rocky Hill, NJ). Nanoparticle formulations were administered to the cells for 24 hours. Supernatants were harvested. Cytokine levels (IL-6, IL-1β, TNF-α) were measured using a Luminex assay (Bio-Techne, Minneapolis, MN).

### Statistical analysis

Statistical analysis was performed using GraphPad Prism software. To determine statistical differences between two groups with normal Gaussian distributions, a Student’s t-test (two-tailed, unpaired, unequal variance, p<0.05) was performed. To determine if statistical differences were significant between three or more groups, one-way ANOVA was performed followed by a post-hoc Tukey’s test. Outliers were identified and removed using Grubbs’ test (p=0.1).

## RESULTS

### Synthesis and characterization of cationic nanoparticles to evaluate mechanisms governing *in vivo* toxicity

We investigated the *in vivo* immune response to bPEI and chitosan nanoparticles and determine how IAA modification of the nanoparticles would alter the immune response. Each nanoparticle formulation was synthesized through an ionic gelation process with sodium tripolyphosphate (unmodified bPEI nanoparticles = bPEI-TPP, IAA-modified bPEI nanoparticles = bPEI-IAA-TPP, unmodified chitosan nanoparticles = Ch-TPP, IAA-modified chitosan nanoparticles = Ch-IAA-TPP; Fig. 1B).[24] Both PEI and chitosan have been reported to activate TLR4 in macrophages, leading to the secretion of IL-12, nitric oxide, and tumor necrosis factor alpha (TNF-α).[25–27] In addition, chitosan triggers DNA release from mitochondria in vitro, which results in stimulator-of-interferon-gene (STING) pathway-mediated upregulation of antiviral signaling through production of interferons and pro-inflammatory cytokines.[28] We hypothesized that these innate immune reactions could be reduced by modification of the primary amines in these polymers to secondary and tertiary amines – which could, at the same time preserve or enhance their buffering capacity and thus delivery efficacy. To achieve this, we conjugated IAA to bPEI or chitosan using a carbodiimide chemistry (Fig. 1C).[11] Using nuclear magnetic resonance spectroscopy (NMR), we determined that 12% of available reaction sites on the bPEI polymer were modified with IAA (Supplementary Fig. 1A). Nanoparticles composed of unmodified or IAA-modified bPEI were slightly less than 100 nm in size (Supplementary Fig. 1B). Both unmodified and IAA-modified bPEI nanoparticles had a similar, slightly negative zeta potential (~-10 mV) at pH 7.4. To quantify IAA modification of chitosan, we measured absorbance by spectrophotometry at 230 nm (Supplementary Fig. 2A). Unmodified chitosan nanoparticles had a mean diameter just below 100 nm. The IAA-modified chitosan nanoparticles had a slightly wider size distribution than their unmodified counterparts,but SEM images suggest that this is due to aggregation, not the formation of larger nanoparticles (Supplementary Fig. 2B-C). As with the bPEI nanoparticles, IAA modification did not affect the zeta potential of the chitosan nanoparticles; both formulations had a slightly positive zeta potential (~+15 mV) at pH 7.4.

**Figure 1.**
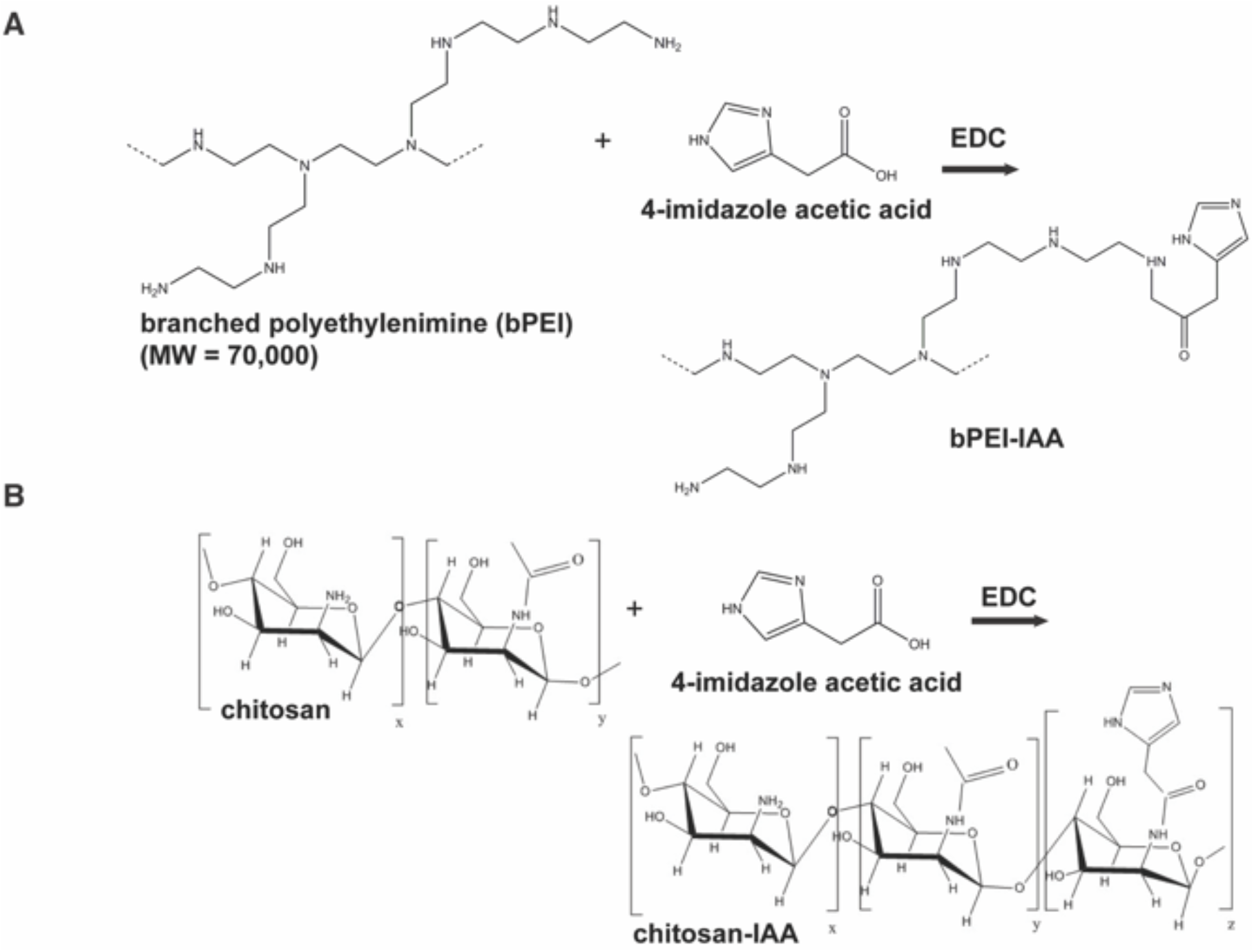
Reaction scheme of imidazole-modified polymers to synthesize nanoparticles for RNA delivery. **(A)** Synthesis of IAA-modified bPEI. **(B)** Synthesis of IAA-modified chitosan.

### IAA modification reduces the *in vivo* toxicity of bPEI and chitosan nanoparticles

We evaluated the systemic toxicity of cationic nanoparticles after intravenous (IV) delivery, which is an administration route associated with high levels of toxicity.[29] Our initial *in vivo* experiments probed serum levels of the pro-inflammatory cytokines IL-6 (Fig. 2A), TNF-α (Fig. 2B), IL-1β (Fig. 2C), and alanine aminotransferase (ALT – a biomarker of liver toxicity; Fig. 2D) 2 hours after nanoparticle administration. Unmodified bPEI nanoparticles elevated levels of IL-6 and ALT in the serum two times more than IAA-modified bPEI nanoparticles did. No detectable amounts of TNF-α and IL-1β were measured after *in vivo* administration of either bPEI formulation. Unmodified chitosan nanoparticles induced *in vivo* production of IL-6, TNF-α, and IL-1β. Behavioral evaluation of mice 30 minutes after IV injection supported our serum cytokine and ALT data. When treated with unmodified chitosan or bPEI nanoparticles, the mice suffered from discomfort and lethargy in comparison to naïve, untreated mice. In mice treated with IAA-modified chitosan or bPEI nanoparticles, the symptoms of lethargy were reduced.

**Figure 2.**
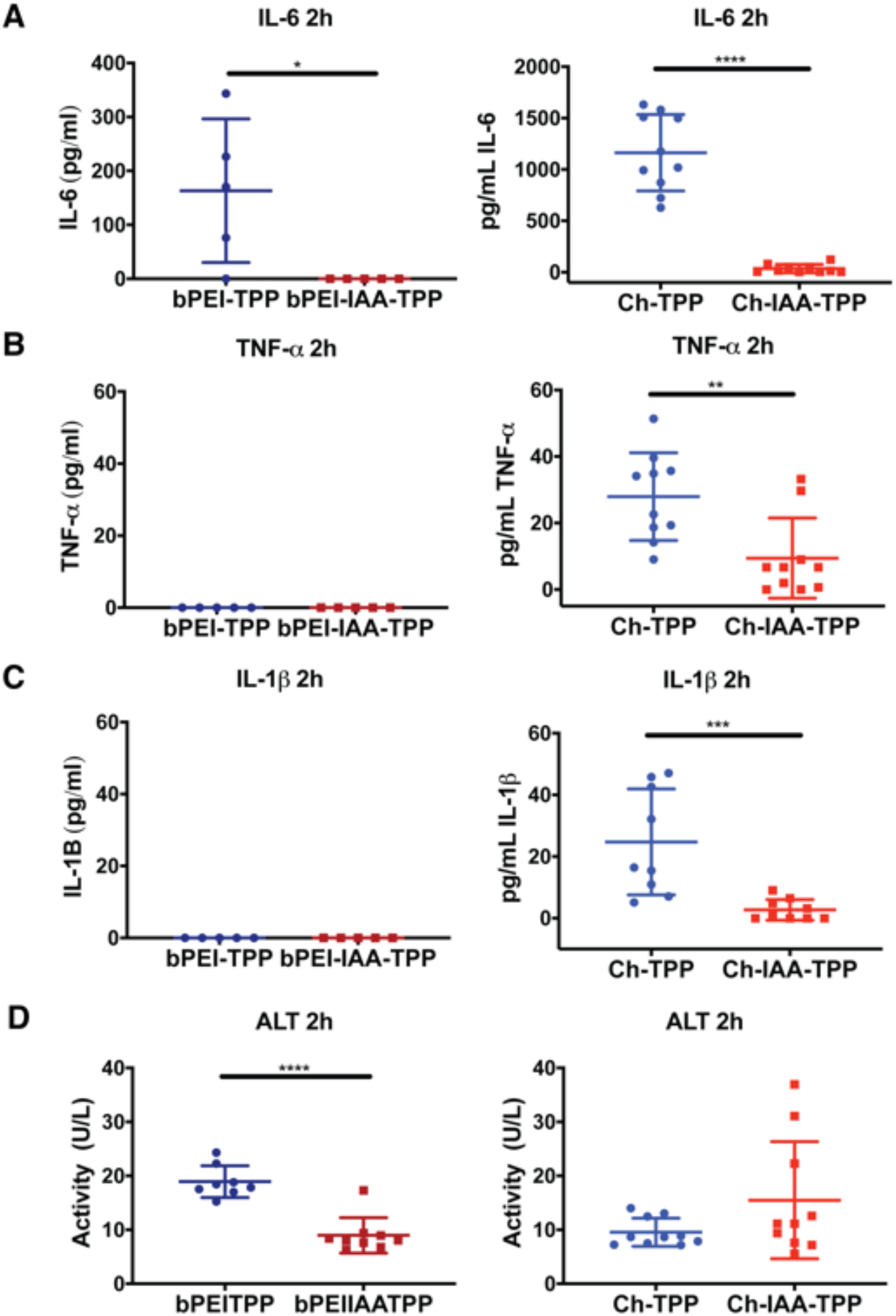
IAA modification reduces in vivo systemic toxicity to bPEI and chitosan nanoparticles. Levels of **(A)** IL-6, **(B)** TNF-**α**, and **(C)** IL-1**β** in serum 2 hours after IV injection of bPEI nanoparticles (unmodified bPEI nanoparticles: bPEI-TPP; IAA-modified bPEI nanoparticles: bPEI-IAA-TPP) or chitosan nanoparticles (unmodified chitosan nanoparticles: Ch-TPP; IAA-modified chitosan nanoparticles: Ch-IAA-TPP; n=5-6). **(D)** Levels of ALT in serum 2 hours after bPEI or chitosan nanoparticle injection (n=8-10). Statistical differences in experiments between two groups were determined using a 2-tailed, unpaired Student’s t-test assuming unequal variance. **P≤ 0.01, \*\*\***P≤0.001, ****P≤ 0.0001**.

Next, we evaluated how IAA modification affects bPEI nanoparticle organ biodistribution, immune cell activation in the spleen, and complement activation. Unmodified, fluorescently labeled bPEI nanoparticles accumulated in the liver, kidney, and lungs one hour after IV injection. The unmodified bPEI nanoparticles remained in these organs 24 hours after injection. IAA-modified, fluorescently labeled bPEI nanoparticles localized in the same organs, but fewer nanoparticles were present in organs 24 hours after injection (Supplementary Fig. 3A-C). In the spleen, unmodified bPEI nanoparticles increased levels of the chemokines (C-X-C motif) ligand 1 (CXCL1), (C-X-C motif) ligand 2 (CXCL2), and (C-C motif) ligand 2 (CCL2) in the spleen (Fig. 3A, Supplementary Fig. 4A). Circulating monocyte uptake of unmodified or IAA-modified bPEI nanoparticles was not significantly different one hour after injection (Fig. 3B). Circulating neutrophil uptake of unmodified bPEI nanoparticles, however, was found to be 3 times higher than uptake of IAA-modified bPEI nanoparticles (Fig. 3C). Complement C3 and C5 activation, measured by complement C3a and C5a levels in the serum, were not significantly different in mice treated with unmodified or IAA-modified bPEI nanoparticles (Fig. 3D-E).

**Figure 3.**
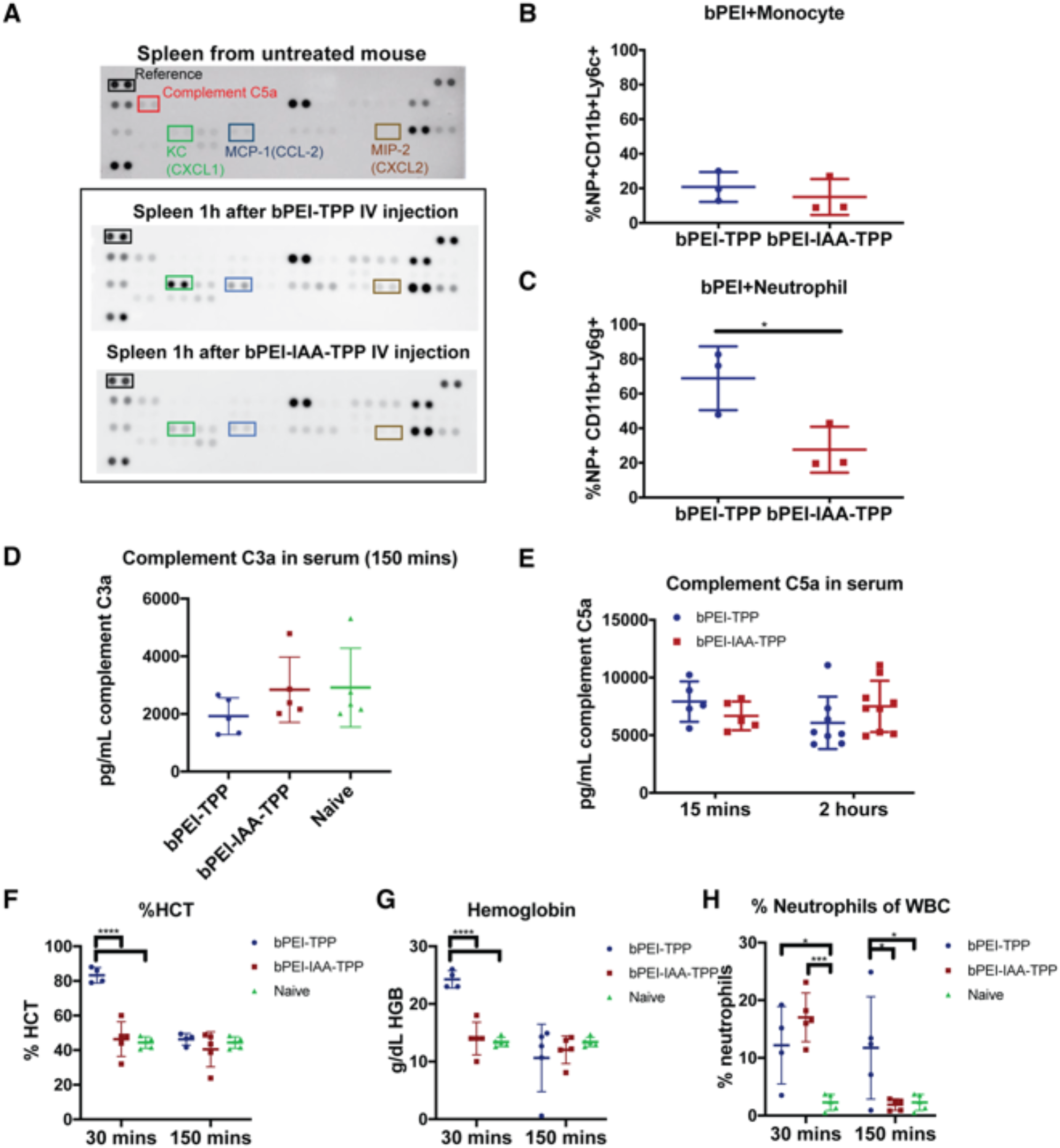
Reduced systemic toxicity of bPEI formulations by IAA modification is dominated by reduced chemokine secretion and the prevention of changes in blood properties. **(A)** Cytokine and chemokine levels in the spleen in a naïve mouse and 1h after IV administration of unmodified bPEI nanoparticles (bPEI-TPP) or IAA-modified bPEI nanoparticles (bPEI-IAA-TPP). Each blot represents one mouse with spots in duplicate and is representative of blots from three treated mice. Representative immunoblots highlighting differences in CXCL1, CCL2, and CXCL2 levels in the spleen after nanoparticle treatment. **(B, C)** The accumulation of bPEI nanoparticles in monocytes and neutrophils in blood 1 hour after IV injection. **(D)** Complement C3a and **€** Complement C5a activation in mice after injection with bPEI nanoparticles. **(F)** Hematocrit levels (HCT), **(G)** hemoglobin concentration (HGB), and the **(H)** percentage of neutrophils of the white blood cell population were measured in whole blood 30 and 150 minutes after bPEI nanoparticle injection (n=4-5). Outliers were identified and removed using the Grubbs test (p=0.1). Error bars represent SD of the mean. Statistical differences in experiments between two groups were determined using a 2-tailed, unpaired Student’s t-test assuming unequal variance. Statistical differences in experiments between more than two groups were determined using one-way ANOVA followed by Tukey’s test. **\*P≤0.05, ***P≤0.001, ****P≤ 0.0001**.

When we collected blood to measure levels of serum cytokines, we observed, qualitatively, differences in blood viscosity in mice treated with the unmodified bPEI nanoparticles. This inspired us to evaluate if cationic nanoparticles influence blood properties after IV injection. 30 minutes after mice were treated with unmodified bPEI nanoparticles, hematocrit rose from 40% to 80%. At the same time point, hemoglobin concentration rose from 12 to 25 g/dL. In contrast, no changes in hematocrit or hemoglobin concentration were observed in IAA-modified bPEI nanoparticle-treated mice after 30 minutes (Fig. 3F-G). Both unmodified and IAA-modified bPEI nanoparticles increased the neutrophil fraction in white blood cells from 0 to 15% after 30 minutes. This neutrophil fraction remained high with mice treated with unmodified bPEI nanoparticles after 150 minutes, but returned to baseline in mice treated with IAA-modified bPEI nanoparticles (Fig. 3H). The increase in hematocrit prompted us to evaluate if the unmodified bPEI nanoparticles were initiating a coagulation cascade by hemolyzing red blood cells, which can induce adenosine diphosphate release and subsequently activate platelets.[30] Neither bPEI nanoparticle formulation induced a significant amount of hemolysis when incubated with whole blood at a concentration of 125 μg/mL, which is twice as high as the blood pool concentration of bPEI nanoparticles immediately after IV injection in our experiments (60 μg/mL; Supplementary Fig. 4B). We then hypothesized that high hematocrit could be attributed to fluid extravasation resulting from enhanced vascular permeability following an anaphylactic response. Histamine levels, however, were not significantly elevated 15 minutes after bPEI nanoparticle injection (Supplementary Fig. 4C) – which suggests a minimal role in anaphylaxis associated with the bPEI nanoparticles. Finally, we assessed the role of TLR4 activation on *in vivo* toxicity by measuring the immune response to bPEI nanoparticles in TLR4 knockout mice. No measurable IL-6 production was measured in the sera of TLR4 knockout mice after 1 hour (Supplementary Fig. 4D), but ALT levels were still elevated (Supplementary Fig. 4E). This indicates that while TLR4 plays a role in the *in vivo* systemic toxicity of bPEI nanoparticles, it is not the sole contributor to its toxic effects.

We performed the same in-depth analysis of *in vivo* biodistribution and systemic toxicity for unmodified and IAA-modified chitosan nanoparticles. The IAA-modified chitosan nanoparticles were more readily internalized by the liver, kidney, and lungs after 1 hour. Moreover, the nanoparticles remained inside these organs 24 hours after administration (Supplementary Fig. 5A-B). Although the IAA-modified chitosan nanoparticles persisted for longer in the mice, we continued to see evidence of reduced *in vivo* systemic toxicity with the IAA-modified chitosan. For instance, the IAA modification prevented the upregulation of CXCL1, CXCL2, and CCL2 observed in the spleens of mice treated with unmodified chitosan nanoparticles after 1 hour (Fig. 4A, Supplementary Fig. 6). Unmodified chitosan nanoparticles were ingested by 60% of circulating monocytes, while IAA-modified chitosan nanoparticles were only ingested by 15% of circulating monocytes (Fig. 4B). Curiously, the circulating neutrophil uptake of chitosan nanoparticles were in agreement with the *in vivo* organ biodistribution data. 65% of circulating neutrophils ingested unmodified chitosan nanoparticles, while nearly 100% of circulating neutrophils ingested IAA-modified chitosan nanoparticles (Fig. 4C). Complement C3a and C5a levels in mice treated with IAA-modified chitosan nanoparticles were nearly double the levels in mice treated with unmodified chitosan nanoparticles (Fig. 4D-E). The unmodified chitosan nanoparticles did not induce higher levels of hematocrit and hemoglobin (Fig. 4F-G), but both unmodified and IAA-modified chitosan nanoparticles increased neutrophil levels in the blood after 30 minutes (Fig. 4H).

**Figure 4.**
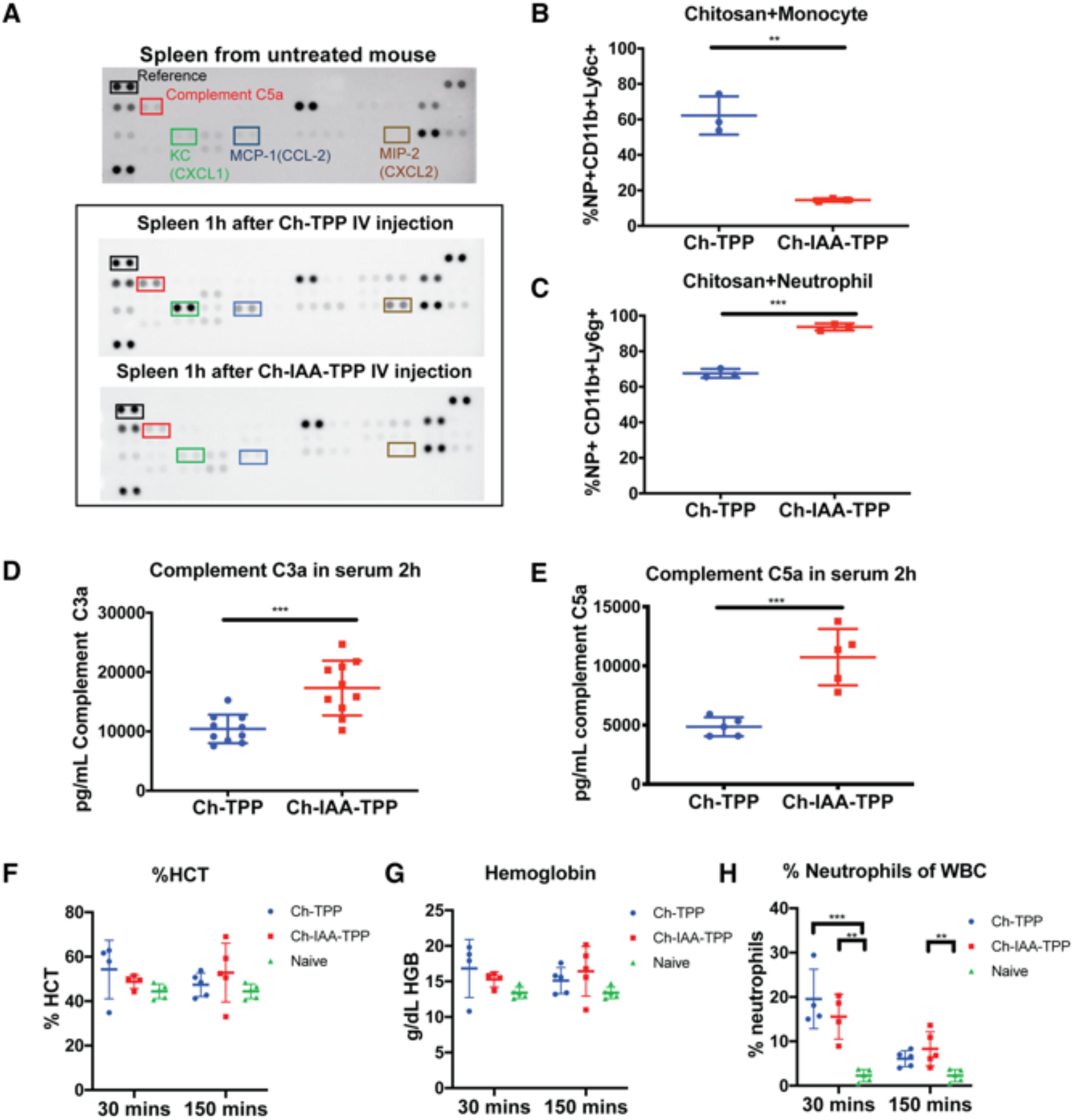
Reduced systemic toxicity of chitosan formulations by IAA modification is driven by complement activation. **(A)** Cytokine and chemokine levels in the spleen in a naïve mouse and 1h after IV administration of unmodified chitosan nanoparticles (Ch-TPP) or IAA-modified chitosan nanoparticles (Ch-IAA-TPP). Each blot represents one mouse with spots in duplicate and is representative of blots from three treated mice. Representative immunoblots highlighting differences in complement C5a, CXCL1, CCL2, and CXCL2 levels in the spleen after nanoparticle treatment. **(B, C)** The accumulation of chitosan nanoparticles in monocytes and neutrophils in blood 1 hour after IV injection. **(D)** Complement C3a and **(E)** Complement C5a activation in mice after injection with chitosan nanoparticles. **F)** Hematocrit levels (HCT), **(G)** hemoglobin concentration (HGB), and the **(H)** percentage of neutrophils of the white blood cell population were measured in whole blood 30 and 150 minutes after nanoparticle injection (n=4-5). Statistical differences in experiments between two groups were determined using a 2-tailed, unpaired Student’s t-test assuming unequal variance. Statistical differences in experiments between more than two groups were determined using one-way ANOVA followed by Tukey’s test. **\*\* P≤0.01, ***P≤0.001**.

### IAA modification significantly changes the protein corona of cationic nanoparticles

We then investigated how the surfaces of the unmodified and IAA-modified nanoparticles changed after exposure to serum proteins *ex vivo*. Mass spectrometry-based proteomic analysis revealed that IAA modification of bPEI and chitosan significantly decreased complement adsorption that typically occurs after incubation in mouse plasma for one hour (Supplementary Fig. 7A-C). Exclusively with the bPEI nanoparticles, IAA modification significantly decreased the adsorption of fibrinogen proteins to the nanoparticle surface (Supplementary Fig. 7D-F). These data suggest that the evolution of the protein corona is dependent on both the type of polymer (e.g., chitosan, bPEI) and if the polymer is modified with IAA.

### *In vitro* screening of unmodified and IAA-modified cationic nanoparticles does not predict their *in vivo* systemic toxicity

To determine if *in vivo* systemic toxicity could be predicted with *in vitro* assays, we evaluated the innate immune response to unmodified and IAA-modified bPEI nanoparticles with mouse bone marrow-derived dendritic cells (BMDCs). We hypothesized that IAA modification reduced systemic toxicity by reducing interactions with TLR4, so we performed a proximity ligation assay to measure the interaction of TLR4 and toll-interleukin 1 receptor domain containing adaptor protein (TIRAP), which is essential for TLR4 signaling to induce pro-inflammatory activation.[31] Indeed, unmodified bPEI nanoparticles triggered TLR4 signaling after 30 minutes, while the IAA-modified bPEI nanoparticles failed to trigger TLR4 signaling (Fig. 5A). Treatment of BMDCs with unmodified bPEI nanoparticles resulted in 2-fold higher secretion of the pro-inflammatory cytokines IL-6 and IL-1β over BMDCs treated with IAA-modified bPEI nanoparticles (Fig. 5B-C). No significant differences in TNF-α secretion were observed between cells treated with unmodified or IAA-modified bPEI nanoparticles (Fig. 5D). Endosomal escape of unmodified and IAA-modified bPEI nanoparticles was evaluated by measurement of their co-localization with intracellular compartments (CD63, EEA1 – an early endosomal marker, LAMP1 – a lysosomal marker, clathrin, and caveolin) after 2 hours of treatment. Incomplete overlap with intracellular compartments was observed with both unmodified and IAA-modified nanoparticle formulations (Supplementary Fig. 8A). This indicates that bPEI nanoparticles, regardless of IAA modification, are able to escape into the cytoplasm to facilitate delivery of mRNA or siRNA. We confirmed that the changes in innate immune activation were not due to differences in cellular uptake of the bPEI and IAA-modified bPEI nanoparticles (Supplementary Fig. 8B).

**Figure 5.**
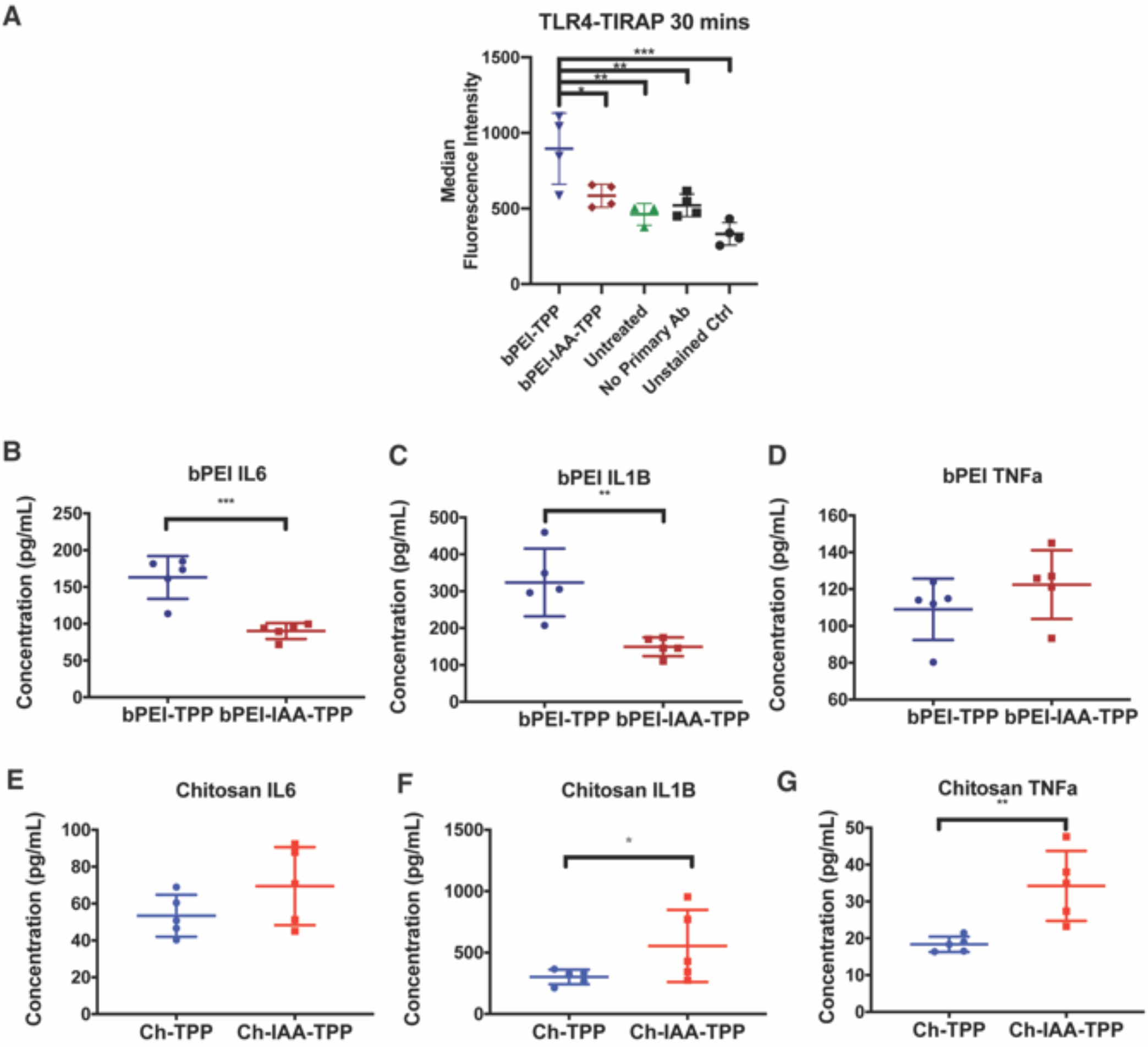
In vitro screening does not predict the in vivo immune response to all cationic nanoparticles. **(A)** Measurement of TLR4-TIRAP interactions 30 minutes after treatment with bPEI nanoparticles in RAW 264.7 macrophages using a flow cytometry-based proximity ligation assay (n=3-4). Outliers were identified and removed using Grubbs’ test (p=0.1). **(B-G)** Mouse BMDCs were treated with unmodified bPEI nanoparticles (bPEI-TPP), IAA-modified bPEI nanoparticles (bPEI-IAA-TPP), unmodified chitosan nanoparticles (Ch-TPP), or IAA-modified chitosan nanoparticles (Ch-IAA-TPP) for 24h. Cell supernatants were collected and IL-6, IL-1**β** and TNF-**α** were measured (n=5). Error bars represent SD of the mean. Statistical differences in experiments between two groups were determined using a 2-tailed, unpaired Student’s t-test assuming unequal variance. Statistical differences in experiments between more than two groups were determined using one-way ANOVA followed by Tukey’s test. **\*P≤0.05, **P≤ 0.01, ***P≤0.001, ****P≤ 0.0001**.

We then tested if IAA modification could reduce the *in vitro* innate immune response to chitosan nanoparticles. First, we evaluated the uptake of unmodified and IAA-modified chitosan nanoparticles by mouse endothelial cells (MEC, Supplementary Fig. 8C). A stark difference was observed in chitosan nanoparticle uptake after IAA modification. We found that 80% of the MECs had internalized IAA-modified chitosan nanoparticles after 24 hours, while fewer than 20% of MECs internalized unmodified chitosan nanoparticles at the same time point. In association with higher cellular uptake, the IAA-modified chitosan nanoparticles were more inflammatory to mouse BMDCs than unmodified chitosan nanoparticles. While there were no significant differences in levels of secreted IL-6 (Fig. 5E), both secreted IL-1β (Fig. 5F) and TNF-α (Fig. 5G) levels were elevated in BMDCs treated with IAA-modified chitosan nanoparticles.

### IAA-modified cationic nanoparticles transfect siRNA and mRNA

With the end goal of designing cationic nanoparticles for nucleic acid transfection, we tested if IAA-modified bPEI and chitosan nanoparticles could transfect siRNA. The PD-L1 gene was targeted in B16-F10 melanoma cells, which is an intriguing therapeutic target that could potentially reduce tumor-related immune suppression.[32] PD-L1 was first induced in melanoma cells through treatment with interferon-γ at a concentration of 50 ng/mL. Treatment of PD-L1-induced cells with unmodified bPEI or chitosan nanoparticles yielded two observations. First, nanoparticles with PD-L1 target siRNA could reduce PD-L1 gene expression relative to nanoparticles with negative control siRNA. Second, unmodified bPEI and chitosan nanoparticles with negative control siRNA upregulated PD-L1 expression over PD-L1-induced cells without nanoparticles. IAA-modified nanoparticles with negative control siRNA, on the other hand, did not upregulate PD-L1 expression. Since PD-L1 can be upregulated in response to TLR4 activation, these data further support that IAA modification reduces the inflammatory response to cationic nanoparticles (Supplementary Fig. 9A-D).[33]

We also evaluated if bPEI nanoparticles could deliver functional luciferase mRNA (mLuc). First, we delivered mLuc with bPEI nanoparticles to HELA cells and measured luminescence from cell lysates after 5 hours. Normalized luminescence signal from IAA-modified bPEI nanoparticle-treated cells with mLuc was four times higher than from cells treated with unmodified bPEI nanoparticles with mLuc (Supplementary Fig. 10A). Following this study, we evaluated if IAA-modified bPEI nanoparticles could deliver functional mLuc in vivo. Three hours after IV delivery of luciferase mRNA, we observed a 20-fold increase in BLI signal in the lungs relative to untreated mice. This level of BLI signal remained high after 6 hours (Supplementary Fig. 10B). The equivalent dose of unmodified bPEI nanoparticles caused severe acute toxicity within 30 minutes, so luciferase expression was not measured.

## DISCUSSION

The primary goal of our study was to identify the principal drivers of *in vivo* systemic toxicity induced by two commonly studied cationic polymer nanoparticles – branched polyethylenimine and chitosan (Table 1) – and evaluate if and how the conversion of primary amines to tertiary amines by modification with IAA could mitigate their *in vivo* systemic toxicity.

**Table 1.**

| Measure of *In Vivo*<br>Immune Response | bPEI | Effect of<br>IAA | Chitosan | Effect of<br>IAA |
| --- | --- | --- | --- | --- |
| **After 30 minutes** |  |  |  |  |
| CBC %hematocrit | ++ | ↓ | + | Same as<br>unmodified |
| CBC hemoglobin | ++ | ↓ | + | Same as<br>unmodified |
| CBC neutrophils | ++ | Same as<br>unmodified | ++ | Same as<br>unmodified |
| **After 1 hour** |  |  |  |  |
| Monocyte uptake | + | Same as<br>unmodified | ++ | ↓ |
| Neutrophil uptake | + | ↓ | + | ↑ |
| Spleen CXCL1 | ++ | ↓ | ++ | ↓ |
| Spleen CCL2 | + | ↓ | + | ↓ |
| Spleen CXCL2 | + | ↓ | ++ | ↓ |
| Spleen Complement C5a | No effect | No effect | + | ↓ |
| **After 2 hours** |  |  |  |  |
| Serum IL-6 | + | ↓ | ++ | ↓ |
| Serum TNF-α | No effect | No effect | + | ↓ |
| Serum IL-1β | No effect | No effect | + | ↓ |
| Serum ALT | ++ | ↓ | No effect | No effect |
| Complement C3 activation<br>(C3a) | No effect | No effect | + | ↑ |
| Complement C5 activation<br>(C5a) | No effect | No effect | + | ↑ |
| CBC Neutrophils (150<br>mins) | + | ↓ | + | Same as<br>unmodified |

Prior reports have evaluated nanoparticle toxicity by measuring *in vitro* innate immune activation in some *ex vivo* immune cells[26, 27, 34–36], *ex vivo* complement activation in serum [30, 37, 38], and changes in the protein corona after *ex vivo* incubation with serum [39–41] There are also a handful of studies that has reported that cationic polymers induce *in vivo* innate immune responses. However, comprehensive evaluation and relative contributions of hematological, hepatic, and immune reactions to the overall systemic toxicity profile of cationic polymer nanoparticles, and mechanism to alleviate them, has not been studied..[25, 28] Although researchers have evaluated the effect of various small molecule modifications on the *in vitro* cytotoxicity of cationic nanoparticles, their effects on *in vivo* systemic toxicity, are poorly understood. Our results address several of these significant gaps in knowledge using chitosan and bPEI as exemplar cationic polymers, and IAA as an exemplar modification that reduces the primary amine content and increases tertiary amines in these polymers.

We discovered that IAA modification could indeed reduce the acute systemic toxicity response to bPEI nanoparticles. *In vivo* systemic toxicity induced by unmodified bPEI nanoparticles is dominated by a combination of changes in blood properties, stimulation of CXCL1 production in the spleen, and liver toxicity measured by increases in ALT levels in the serum (Fig. 6A). To our knowledge, this is the first report identifying cationic polymer nanoparticle-induced changes to blood properties *in vivo*. We infer that the sharp increase in hematocrit after treatment with unmodified bPEI nanoparticles is associated with liver toxicity, as studies have correlated heightened levels of ALT to high hematocrit.[24, 42] IL-6 has been shown to increase production of acute phase proteins and CXCL1, which suggests that the changes in blood properties after bPEI nanoparticle administration could be a consequence of hepatocyte damage.[43] CXCL1 can then recruit neutrophils into circulation and aid in the clearing of unmodified bPEI nanoparticles.[44] Our discovery of heightened liver toxicity is in line with reports that demonstrate the expression of TLR4 on hepatocytes, elevated hepatocyte toxicity to PEI-coated gold nanoparticles, and PEI-induced impairment of cellular respiration in liver mitochondria.[34, 41, 45] However, our results on systemic toxicity in TLR4 knockout mice, shows that TLR4 is not the only driver of hepatic toxicity to bPEI nanoparticles. This conclusion is based on the elimination of the IL-6 response, but not the ALT response after injection with unmodified bPEI nanoparticles in TLR4 knockout mice. Remarkably, IAA was able to downregulate several biomarkers of *in vivo* systemic toxicity to bPEI nanoparticles. The IAA modification reduced liver toxicity, IL-6 production, and CXCL1 production after intravenous administration. Reduced toxicity may be attributed to changes in the protein corona, in which fibrinogen binding to PEI nanoparticles was significantly reduced after IAA modification. Fibrinogen can enhance innate immune activation and upregulate chemokine expression[46], which was observed after treatment with unmodified PEI nanoparticles but not IAA-modified PEI nanoparticles. The reduced toxicity could not be attributed to changes in complement activation, blood hemolysis, or anaphylactic responses mediated by histamine.

**Figure 6.**
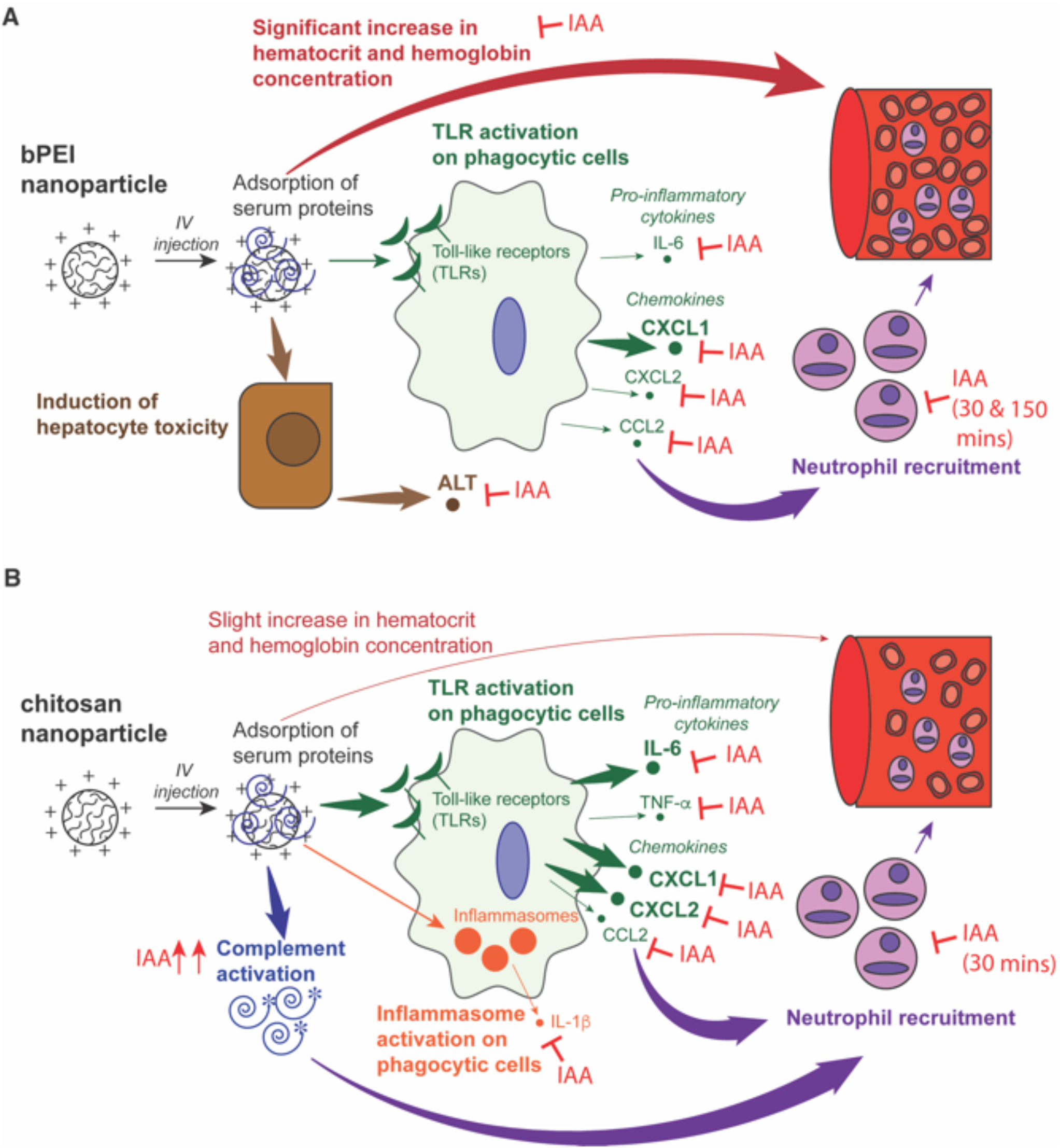
**(A)** In vivo toxicity of bPEI nanoparticles is primarily attributed to a significant increase in hematocrit and hemoglobin concentration, the induction of hepatocyte toxicity, and cytokine and chemokine production by phagocytic cells after TLR activation. **(B)** In vivo toxicity of chitosan nanoparticles is primarily attributed to complement activation, cytokine and chemokine production by phagocytic cells after TLR activation, and inflammasome activation on phagocytic cells.

Modification of chitosan with IAA also reduced its *in vivo* systemic toxicity. This result is fascinating because the *in vivo* systemic toxicity response to unmodified chitosan is not identical to the response to unmodified PEI. The *in vivo* systemic toxicity response of bPEI and chitosan shared elevated neutrophil levels in circulation, IL-6 levels in the serum, and CXCL1 levels in the spleen (Fig. 6B). However, we found that liver toxicity and changes in blood properties do not appear to play a major role in driving *in vivo* systemic toxicity to chitosan. Chitosan nanoparticles also increased serum levels of TNF-α and IL-1β, which is in agreement with studies demonstrating that chitosan activates the inflammasome pathway.[28, 35] [36] IAA modification prevented upregulation of IL-6, CXCL1, and IL-1β by chitosan nanoparticles – which indicate that as with bPEI, IAA reduces innate immune activation by chitosan. At the same time, IAA modification also increased neutrophil uptake of chitosan nanoparticles and *in vivo* complement activation. This seems paradoxical, given that complement activation is typically considered pro-inflammatory. Complement fragments, however, have been shown to enhance the anti-inflammatory activity of neutrophil extracellular traps on TLR4-activated macrophages. [47],[48] Evidence of enhanced neutrophil uptake, heightened complement activation, and reduced IL-6 enable this to be a plausible mechanism for the reduced *in vivo* systemic toxicity of IAA-modified chitosan. This complex phenomenon between multiple cell types could explain the disparity between our *in vivo* and *in vitro* toxicity results with chitosan – where IAA-modified chitosan nanoparticles induced a strong innate immune response from BMDCs. When toxicity is evaluated *in vitro*, the internalization rate of the nanoparticles is the primary driver of toxicity – as chitosan is known to activate the cGAS sensor in the cytosol. Simplified *in vitro* screening assays do not take into consideration that nanoparticles are internalized by a variety of cells *in vivo* or the potential effect of the complement response on toxicity. These *in vitro* screening assays also neglect the effect of acute responses from one cell type that can activate feedback mechanisms in other cell types. In the context of designing nanoparticles for nucleic acid delivery, the mismatch between *in vitro* cytotoxicity and *in vivo* systemic toxicity may falsely eliminate viable nanoparticles from further testing. Therefore, our data demonstrate that thorough *in vivo* toxicity analyses are essential for the advancement of cationic polymer nanoparticles for nucleic acid delivery into the clinic.

## CONCLUSIONS

Taken together, our findings illustrate that *in vivo* systemic toxicity is regulated by a combination of different biological phenomena – TLR activation, complement activation, liver toxicity, and changes in hematology. The contributions of each biological phenomenon to cationic nanomaterial toxicity is different for different materials – for bPEI, toxicity profile is driven by liver toxicity and changes in hematology; while for chitosan, the toxicity profile is governed by a balance of TLR and complement activation. The classification of what makes a “safe nanoparticle” must be determined by a weighted consideration of each of these *in vivo* biological phenomena. We also conclude that chemical modifications do not affect toxicity in the same manner when applied to different polymers. Our results also show that *in vitro* measures of the innate immune response, as with the example of IAA-modified chitosan, do not accurately predict *in vivo* toxicity. Therefore, it is essential to move beyond *in vitro* cytotoxicity screens for nanomaterials to more informative, and detailed *in vivo* toxicity tests to appropriately select nanoparticle candidates for clinical nucleic acid delivery.

## ACKNOWLEDGEMENTS

We thank David Smalley of the Proteomics Core for assistance with proteomic analysis, Jiri Schimer for help with the characterization of the IAA-modified polymers, Wen Seeto and Wesley Grove for help with the CBC analysis, Jared Beyersdorf and Kevin Lindsay for preparing and providing the luciferase mRNA, Cedrick Young and Navdeep Jhita for assistance with animal experiments, Katelynn Phang for assistance with Luminex assays, and Sushma Bhosle for helpful discussions on the design of the *in vitro* transfection assay. Microscopy experiments and flow cytometry experiments were performed in the Engineered Biosystems Building Optical Microscopy Core and Cellular Analysis Core, respectively. Emmeline Blanchard was supported by National Science Foundation Graduate Research Fellowship Program [Grant No. DGE-1650044]. We acknowledge funding support from the Georgia Tech Foundation and the Robert A. Milton Chaired Professorship to Prof. Krishnendu Roy.

## AUTHOR CONTRIBUTIONS

R.T. and K.R. conceived the study. R.T., P.P., V.R., N.D., B.L., J.L, E.B., and P.S. contributed to the experimental design and performed the experiments. R.T., P.P, and N.D. analyzed the data. R.T., D.S., and K.R. wrote the manuscript. All authors edited and reviewed the manuscript.

## DATA AVAILABILITY

The raw data required to reproduce these findings cannot be shared at this time as the data also forms part of an ongoing study. The processed data required to reproduce these findings cannot be shared at this time as the data also forms part of an ongoing study.

