## Supplementary Information for "Modification of primary amines to higher order amines reduces in vivo hematological and immunotoxicity of cationic nanocarriers through TLR4 and complement pathways"

**SUPPLEMENTARY METHODS**

**Synthesis and characterization of bPEI-IAA**

IAA-modified bPEI (bPEI-IAA) was produced by conjugation of branched polyethylenimine (30% solution in water, MW 70,000, Polysciences, Warrington, PA) to imidazole-4-acetic acid (Sigma Aldrich). To synthesize bPEI-IAA, a 1.5% w/v solution of bPEI was prepared by adding 300 mg of bPEI to 20 mL of 0.2 M MES buffer (pH=6.5). A 2% w/v solution of IAA was prepared by adding 84 mg of IAA to 4.2 mL of 0.2 M MES buffer (pH = 6.5). A 2 molar excess of 1-Ethyl-3-(3-dimethylaminopropyl) carbodiimide (EDC, Thermo Fisher Scientific) was then added to the IAA solution. The IAA-EDC mixture was then added to the bPEI solution, vortexed for 1 minute, and reacted overnight on a rotator at room temperature. To remove unreacted EDC, the bPEI-IAA polymer was dialyzed for 36 hours against DI H2O in a Snakeskin membrane (MWCO = 10k). The bPEI-IAA polymer was then lyophilized and resuspended at a concentration of 3 mg/mL in DI H2O for nanoparticle synthesis or in D2O for ^1^H NMR, which was used to quantify the degree of imidazole modification of bPEI-IAA.

**Synthesis and characterization of Ch-IAA**

Chitosan-IAA (Ch-IAA) was produced by conjugation of chitosan PCL113 (Novamatrix, Sandvika, Norway) to imidazole-4-acetic acid (Sigma Aldrich, Saint Louis, MO) as previously reported. To synthesize chitosan-IAA, a 1% w/v solution of chitosan was prepared by adding 100 mg of chitosan to 10 mL of 0.2 M MES buffer (pH=6.5). A 2% w/v solution of IAA was prepared by adding 40 mg of IAA to 2 mL of 0.2 M MES buffer (pH = 6.5). A 20 molar excess of 1-Ethyl-3-(3-dimethylaminopropyl) carbodiimide (EDC) was then added to the IAA solution. The IAA-EDC mixture was then added to the chitosan solution, vortexed for 1 minute, and reacted overnight on a rotator at room temperature. Subsequently, the chitosan-IAA mixture was dialyzed against 5 mM HCl (2x) and deionized water (2x) and lyophilized for 48 hours. IAA modification of chitosan was quantitatively measured through readings of UV absorbance at 230 nm. A standard curve was generated from solutions of IAA of known concentrations in acetate buffer. Sample absorbance was subtracted from the absorbance of an equivalent concentration of chitosan in acetate buffer. Percent modification was calculated with the degree of chitosan acetylation (83%) taken into account.

***In vitro* hemolysis assay**

Whole blood was collected into vials with lithium heparin anti-coagulant. Blood was diluted into PBS (5% w/v) and nanoparticles were added to a concentration of 125 μg/mL. Red blood cell lysis buffer (eBioscience, San Diego, CA) was used as a positive control while PBS was used as a negative control. Samples were incubated for 10 minutes at room temperature. The samples were then centrifuged at 500g for 5 minutes and the supernatants were collected. Each supernatant was diluted 2-fold in PBS and absorbance was measured at 540 nm using a spectrophotometer. Percent hemolysis was calculated by the following equation: (sample absorbance – negative control absorbance)/ (positive control absorbance – negative control absorbance) *100.

***In vivo* nanoparticle organ biodistribution study**

Mice were sacrificed at 1 hour, 6 hours, and 24 hours after IV injection of nanoparticles (8 mg chitosan/kg body weight or 3 mg bPEI/kg body weight). The liver, spleen, lungs, kidneys, and heart were immediately harvested and imaged with an IVIS Spectrum (Perkin Elmer, Waltham, MA). ROIs were drawn in the Living Image software and the radiant efficiency was quantified for each organ. Normalized radiant efficiency was calculated by dividing the radiant efficiency of each organ by the radiant efficiency of an injection volume of the nanoparticle formulation. An injection volume of each nanoparticle formulation was imaged in a well of a 96-well black Costar plate.

**Assessment of nanoparticle protein corona with LC-MS**

Identification of components of the nanoparticle protein corona was performed similar to a published protocol.^33^ Briefly, nanoparticles were formulated and incubated for 1 hour in mouse plasma at 37 C. The total nanoparticle surface area to plasma volume ratio was kept constant between the two formulations. After particle incubation, the nanoparticles in plasma were added to a 0.7 M sucrose cushion and centrifuged at 15,300g. The cushion was removed and the particles were washed three times with 1 mL PBS to remove loosely adhered soft corona proteins. The nanoparticle pellets were resuspended in 1% SDS in 100 mM ammonium bicarbonate and a micro BCA protein assay was performed. For each sample, 40 mg of protein was reduced, alkylated, and digested according to the FASP protocol.^34^ The peptides were analyzed by HPLC-MS/MS. Peptides were separated on 2 micron beads in a 50 cm x 75 μm C18 column and were analyzed with a Q Exactive Plus mass spectrometer. Each individual sample was run twice and conditions were run in triplicate. The spectra were searched against a user provided database using the Mascot Search algorithm via Proteome Discoverer 2.1. A recently downloaded UniProt mouse FASP database was used. Search parameters included variable modifications for oxidation of Met, carboxyamidomethylation of Cys residues, and acetylation of protein N-terminal.

***In vitro*** **evaluation of nanoparticle endosomal escape**

Glass coverslips were sterilized and placed into 24 well plates. HELA cells were seeded on the coverslips at a density of 60,000 cells/well and allowed to grow overnight. Fluorescently labeled nanoparticles were added to the cells and incubated for 2 hours. Cells were washed 3x with PBS and media was replaced. The cells were returned to the incubator for 2 more hours. At this timepoint, cells were washed 3x with PBS and fixed with BD Cytofix buffer for 10 minutes. Cells were washed 3x with PBS and permeabilized with 0.002% Triton X for 10 minutes. The cells were washed 3x with PBS then blocked with 10% donkey serum overnight at 4 C. Following overnight incubation, cells were washed 3x with PBS and incubated with a cocktail of primary antibodies to intracellular compartment markers - clathrin (1:500, Biolegend MMS-423P), caveolin (1:200, Santa Cruz sc-53564), CD63 (1:50, Abcam ab193349), EEA1 (1:100, Santa Cruz sc-365652), and LAMP1 (1:200, Abcam ab25630) – for 30 minutes at 37 C. Cells were washed 3x with PBS and then incubated with an Alexa-Fluor 488 donkey-anti-mouse secondary antibody (1:250, Thermo Fisher) for 30 minutes at 37 C. Next, cells were washed 3x with PBS and the coverslips were mounted onto glass slides using Prolong Gold with DAPI. Slides were imaged on a Perkin Elmer Spinning Disk microscope with an EM-CCD camera (60x magnification). M1 overlap coefficients (nanoparticles to intracellular compartments) were determined using the Co-localization toolbox in Volocity software for individual cells. The percentage of nanoparticles outside of intracellular compartments was calculated as (1 – M1 overlap coefficient) *100.

***In vitro* nanoparticle uptake study**

Cells were seeded in 24 well plates at a density of 60,000 cells/well and allowed to grow overnight. Fluorescently labeled nanoparticles were added to the cells and incubated for 2 hours. Cells were washed 3x with PBS and then incubated with 0.25% trypsin at 37 C for 5 minutes. The cells were then transferred to FACS tubes and 1 mL of FACS buffer (2% FBS in PBS) was added. Samples were washed, fixed with BD Cytofix buffer, washed again, and then resuspended in FACS buffer. Populations of cells with fluorescent particles were measured on the APC channel using a BD Accuri flow cytometer.

**Synthesis and characterization of RNA-loaded bPEI nanoparticles**

Unmodified bPEI nanoparticles and IAA-modified bPEI nanoparticles were produced by mixing solutions of bPEI or bPEI-IAA in deionized water with sodium tripolyphosphate in deionized water. The mass ratio of bPEI or bPEI-IAA to TPP was 3:2. Nanoparticle formulations were then put onto a rotator at room temperature for 30 minutes. Following vortexing, nanoparticles were centrifuged at 4000 g for 20 minutes in Amicon centrifuge filters (100 kD MWCO) and resuspended in 1 mL RNAse free DI H2O. RNA was loaded to nanoparticles at a density of 415 μg RNA/mg bPEI or bPEI-IAA. The nanoparticles were then vortexed for 30 seconds and subsequently placed on an end-to-end rotator at 4 C for 30 minutes. RNA condensation onto the nanoparticles was verified using the Ribogreen assay (Thermo Fisher Scientific).

***In vitro* PD-L1 gene silencing studies**

Negative control siGenome siRNA and PD-L1 siRNA were purchased from Dharmacon (GE Healthcare, UK). Primers (mouse b-actin, PD-L1) and SYBR-Green reagents for RT-PCR were also purchased from Qiagen. B16-F10 cells were cultured on 24-well plates and induced to express PD-L1 through incubation with mouse IFN-γ at a concentration of 50 ng/mL. After 24 hours, nanoparticle formulations with siRNA (50 μg chitosan/well, 7.5 μg bPEI/well) were added at a dose of 1 μg siRNA/target gene. The control formulation was loaded with an equivalent mass of negative control siRNA. After 48 hours of incubation, cells were lysed using the Trizol reagent (Ambion, Thermo Fisher Scientific, Waltham, MA). Total RNA was converted into cDNA using a Superscript (Invitrogen) PCR reaction. qRT-PCR was conducted on an ABI Step-One-Plus real-time PCR system (Applied Biosystems, Foster City, CA) in 12 μl reactions. Each sample was analyzed in duplicate for both target and endogenous control (mouse beta-actin). Fold change in gene expression was assessed using the ΔΔC_t_ quantitation method.

***In vitro* luciferase transfection studies**

Luciferase mRNA was prepared by the lab of Philip Santangelo. HELA cells were seeded in 24 well plates at a density of 60,000 cells/well and allowed to grow overnight. Nanoparticles (1.2 μg bPEI/well) loaded with 500 ng luciferase mRNA were added to each well. At 5 hours, cells were washed three times with PBS. Cell lysate was prepared using the ONE-Glo Luciferase Assay System (Promega). Luminescence was then measured using a BIOTEK plate reader (Gain = 135, 1 second integration time). Measurements were normalized to protein concentration measured by a BCA assay (Thermo Fisher Scientific).

***In vivo* luciferase transfection and bioluminescence imaging**

Mice were injected IV with 60 μg bPEI nanoparticles to deliver a 1.25 mg/kg dose of luciferase mRNA. Prior to imaging, hair was removed from the abdomen of the mice using Nair. An IVIS Spectrum was used to acquire bioluminescence images at 1 hour and 5 hours after injection (exposure time = 1 minute). A 150 mg/kg dose of firefly D-luciferin potassium salt (GoldBio) was injected intraperitoneally 10 minutes before each imaging session.

**SUPPLEMENTARY FIGURES**

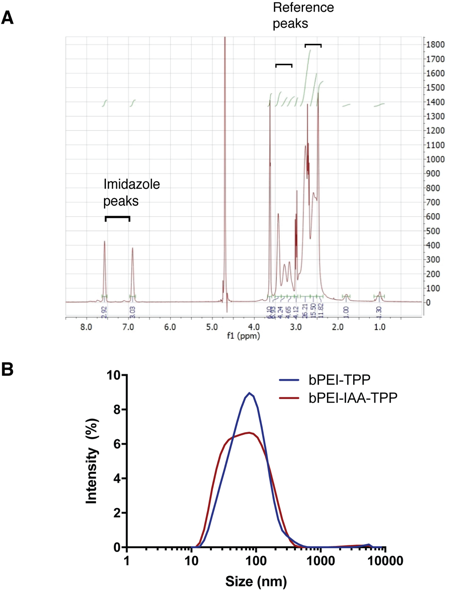

**Supplementary Figure 1. Synthesis and characterization of imidazole-modified PEI (bPEI-IAA) polymer and nanoparticles**. **(A)** NMR spectra for bPEI-IAA. **(B,C)** Size distribution of unmodified bPEI nanoparticles (bPEI-TPP) and IAA-modified bPEI nanoparticles (bPEI-IAA-TPP).

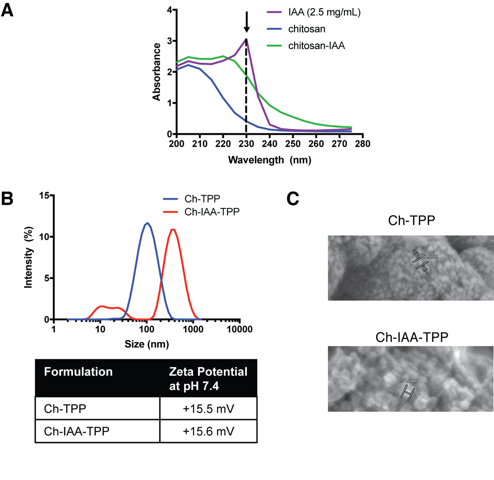

**Supplementary Figure 2. Synthesis and characterization of imidazole-modified chitosan (Ch-IAA) polymer and nanoparticles**. **(A)** Quantitative method to measure level of imidazole modification of chitosan by reading absorbance at 230 nm. **(B)** Size distribution measured by dynamic light scattering and zeta potential of unmodified chitosan and IAA-modified chitosan nanoparticles (Ch-TPP and Ch-IAA-TPP, respectively) **(C)** SEM images of Ch-TPP (60,100x) and Ch-IAA-TPP (45,000x) nanoparticles.

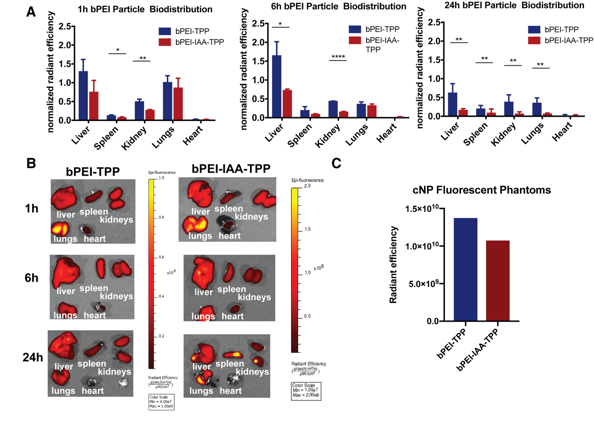

**Supplementary Figure 3. Biodistribution of unmodified bPEI nanoparticles (bPEI-TPP) and IAA-modified bPEI nanoparticles (bPEI-IAA-TPP).**  **(A), (B)** The nanoparticles were labeled with Vivotag-645 and injected IV into wild-type mice. IVIS images of nanoparticle biodistribution in organs were acquired 1 hour, 6 hours, and 24 hours post-injection. Radiant efficiencies of nanoparticles in liver, spleen, kidney, lungs, and heart were normalized to the fluorescence of the injected dose (n=3-4 animals). (**C)** Fluorescence of labeled nanoparticle formulations as measured by IVIS**.** Error bars represent SD of the mean. Statistical differences between groups were determined using a 2-tailed, unpaired Student’s t-test assuming unequal variance. *P≤0.05, **P≤0.01, ****P≤ 0.0001.

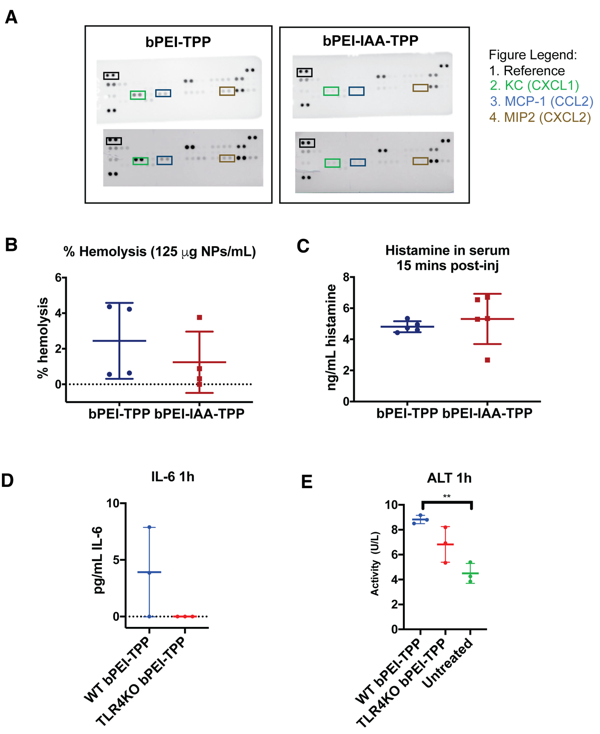

**Supplementary Figure 4.** **Mechanism of reduced immunotoxicity of bPEI nanoparticles by IAA modification. (A)** Immunoblots of spleen 1h after IV administration of unmodified bPEI (bPEI-TPP) IAA-modified bPEI (bPEI-IAA-TPP) nanoparticles. **(B)** The percentage of hemolyzed red blood cells was measured after co-incubation of whole blood (5% v/v) with bPEI nanoparticles (125 µg/mL, n=4 per group). **(C)** Levels of histamine 15 minutes after bPEI nanoparticle injection (n=5). **(D)** IL-6 levels and **(E)** alanine aminotransferase **(**ALT) levels in serum 1h after IV administration of bPEI-TPP nanoparticles in wild-type (WT) or TLR4 knockout (TLR4 KO) mice. Error bars represent SD of the mean. Statistical differences in experiments between two groups were determined using a 2-tailed, unpaired Student’s t-test assuming unequal variance. Statistical differences in experiments between more than two groups were determined using one-way ANOVA followed by Tukey’s test. **P≤ 0.01.

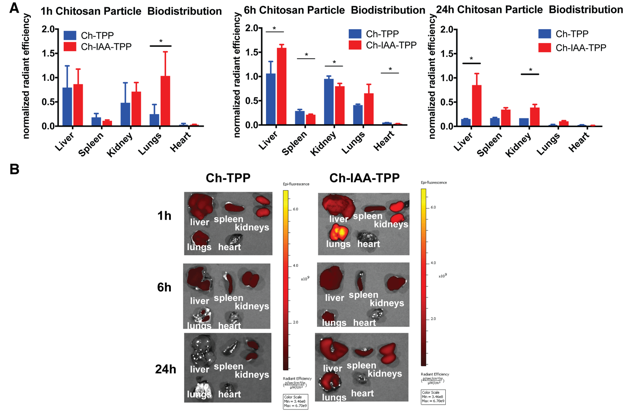

**Supplementary Figure 5. Biodistribution of unmodified chitosan (Ch-TPP) and IAA-modified chitosan nanoparticles (Ch-IAA-TPP).**  **(A), (B)** The nanoparticles were labeled with Vivotag-645 and injected IV into wild-type mice. IVIS images of nanoparticle biodistribution in organs were acquired 1 hour, 6 hours, and 24 hours post-injection. Radiant efficiencies of nanoparticles in liver, spleen, kidney, lungs, and heart were normalized to the fluorescence of the injected dose (n=3-4 animals). Error bars represent SD of the mean. Statistical differences between groups were determined using a 2-tailed, unpaired Student’s t-test assuming unequal variance. *P≤0.05.

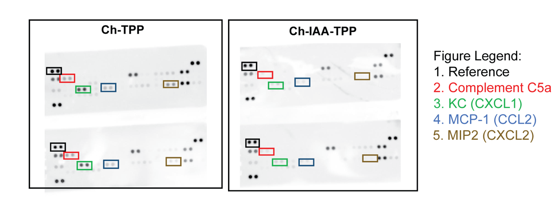

**Supplementary Figure 6.** **Spleen immunoblots from mice treated with chitosan nanoparticles 1h after IV injection.**

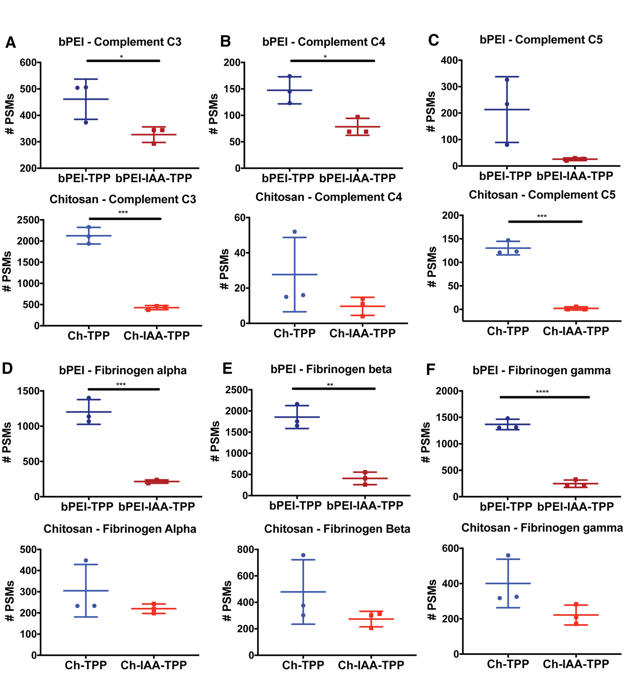

**Supplementary Figure 7. *In vitro* evaluation of the protein corona surrounding bPEI and chitosan nanoparticles after exposure to mouse plasma.** **(A)** Complement C3, **(B)** complement C4, **(C)** complement C5, **(D)** fibrinogen alpha, **(E)** fibrinogen beta, and **(F)** fibrinogen gamma bound to the surface of unmodified or IAA-modified bPEI or chitosan nanoparticles after incubation in mouse plasma for one hour, quantified in peptide spectral matches (PSMs, n=3). Statistical differences in experiments between two groups were determined using a 2-tailed, unpaired Student’s t-test assuming unequal variance. *P≤0.05, **P≤ 0.01, ***P≤0.001, ****P≤ 0.0001.

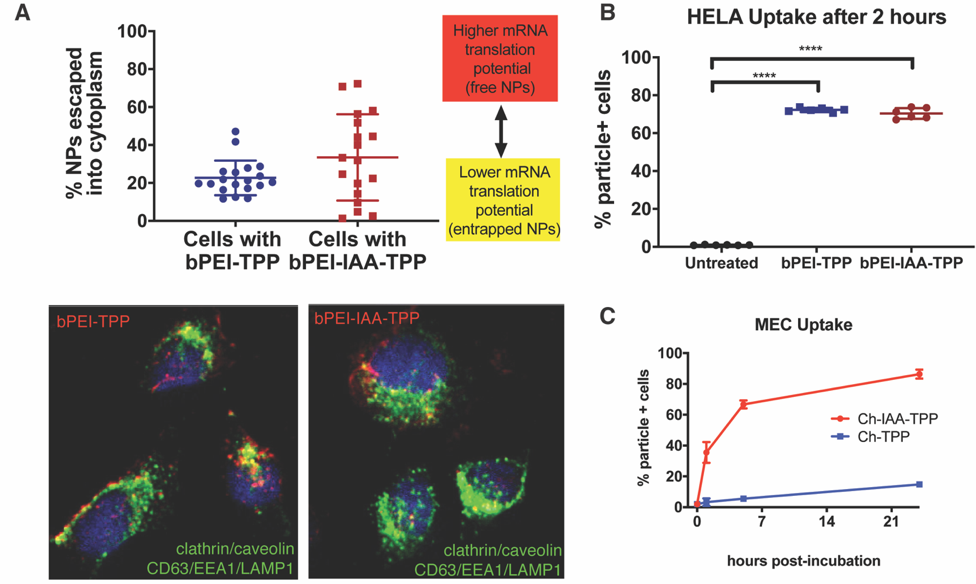

**Supplementary Figure 8. *In vitro* endosomal escape and cell uptake of cationic nanoparticles.** **(A)** Percentage of cells with more than 15% of bPEI or IAA-modified bPEI nanoparticles escaped from intracellular compartments (CD63, EEA1, LAMP1, clathrin, caveolin) in HELA cells after 4 hours (n=18-19 cells). **(B)** In vitro uptake of nanoparticles by HELA cells after 2 hours (n=6). Error bars represent SD of the mean. **(C)** Uptake of unmodified chitosan and IAA-modified chitosan nanoparticles in mouse endothelial cells (MECs). Statistical differences in experiments between two groups were determined using a 2-tailed, unpaired Student’s t-test assuming unequal variance. Statistical differences in experiments between more than two groups were determined using one-way ANOVA followed by Tukey’s test. ****P≤ 0.0001.

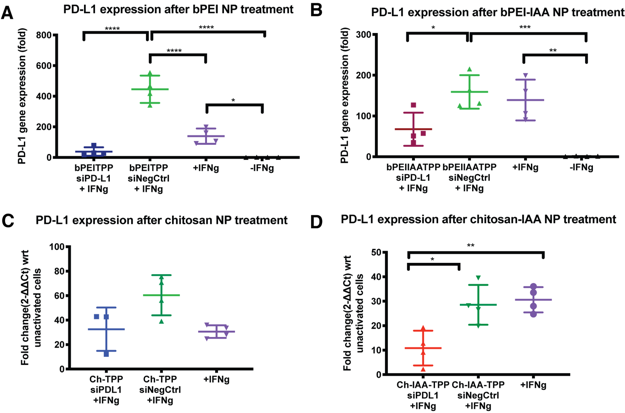

**Supplementary Figure 9. IAA-modified bPEI nanoparticles transfect siRNA *in vitro.*** PD-L1 expression was induced in B16-F10 cells by addition of IFN-𝝲 (50 ng/mL).  PD-L1 induced B16-F10 cells were treated with **(A)** unmodified bPEI nanoparticles (bPEI-TPP), **(B)** IAA-modified bPEI nanoparticles (bPEI-IAA-TPP), **(C)** unmodified chitosan nanoparticles (Ch-TPP), **(D)** or IAA-modified chitosan nanoparticles (Ch-IAA-TPP) loaded with 1 µg PD-L1 siRNA (siPDL1) or negative control siRNA (siNegCtrl, n=3-4).  PD-L1 expression was measured and normalized to cells that were not exposed to IFN-𝝲. Error bars represent SD of the mean. Statistical differences between groups were determined using 1-way ANOVA and Tukey’s test. *P≤0.05, **P≤0.01 ***P≤0.001, ****P≤ 0.0001.

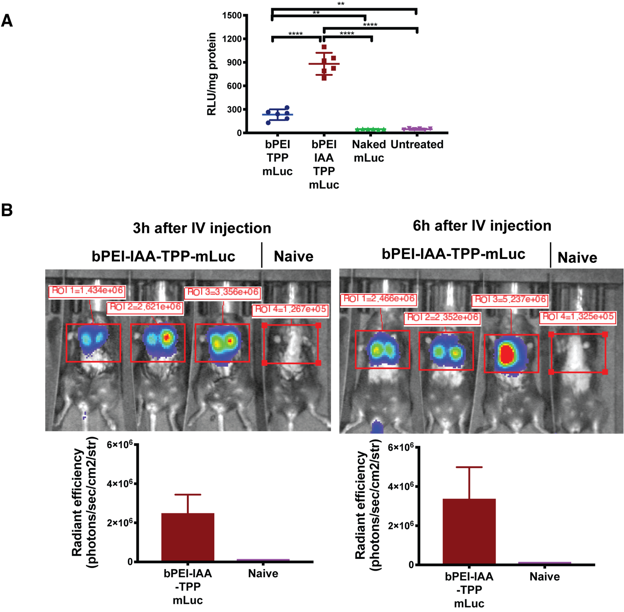

**Supplementary Figure 10. IAA-modified bPEI nanoparticles transfect mRNA in vitro and in vivo. (A)** Luciferase expression in HELA cells 5 hours after treatment with naked mLuc or bPEI nanoparticles loaded with mLuc (500 ng dose, n=6). **(B)** Luciferase expression in mice 3 hours and 6 hours after IV injection of IAA-modified bPEI nanoparticles loaded with 25 µg mLuc. Error bars represent SD of the mean. Statistical significance was measured using one-way ANOVA followed by Tukey’s test for multiple comparisons. *P< 0.05, **P<0.01, ****P<0.0001.
